# Deep2Full: Predictive model for complementing phenotypic outcomes in a deep mutational scan using protein sequence and structure information

**DOI:** 10.1101/217158

**Authors:** C. K. Sruthi, Meher K. Prakash

## Abstract

Large scale mutagenesis experiments are becoming possible owing to the advancement in the sequencing technologies and high throughput screening. Deep mutational scans perform exhaustive single-point mutations on a protein and probe their phenotypic effects. Performing a full scan with site-directed mutations of all the amino acid residues in a protein may not be practical, and may not even be required, especially if predictive computational models can be developed. Computational models are however naive to cellular response in the myriads of assay-conditions. In order to develop the realistic paradigm of assay context-aware predictive hybrid models, we combine minimal deep mutational studies with computational models and predict the phenotypic outcomes quantitatively. Structural, sequence and co-evolutionary information along with partial deep mutational scan data was included to capture the phenotypic relevance of the mutations to the specific screening criterion. The model reliably predicts the fitness outcomes of hundreds of randomly selected amino acid mutations in β-lactamase, when the phenotypic fitness data from as few as 15% of the full mutation is available. Interestingly, the predictive capabilities are better with a random set of mutations rather than with a systematic substitution of all amino acids to alanine, asparagine and histidine (ANH). The model can potentially be extended for predicting the phenotypic outcomes at other concentrations of the stressor by carefully analyzing the dose-response curves of a representative set of mutations.

**Author Summary:** Mutations are the minor changes in protein sequences, with incommensurately high consequences for their function. Many severe diseases can occur with simple single point mutations. An interesting way of studying these mutations is not to isolate the protein from its natural conditions, but rather study how the fitness of the cell improves or decreases in response to these mutations. Whether it is for understanding disease biology or for bio-engineering applications it is important to quantify the impact of mutations on the cellular fitness. An experimental paradigm has evolved which has improved the ability to sample several hundred thousands of mutation-fitness relations using high throughput screening. However, since these are very specialized experiments, the question is if the number of such experiments required can be minimized, by using computer models to complement the rest of the fitness predictions. In this work we introduce this new paradigm which uses computer model trained on a partial deep mutation scan data, to predict the fitness variations in a full mutations scan that could also be repeated under multiple experimental conditions like drug concentrations.

## 1 Introduction

Genetic diversity arises from mutations. Interestingly a very large portion of genetic variation is represented by single nucleotide polymorphisms (SNPs), ^1,2^ which is at most a variation in a single amino acid in a protein. The effect of mutations, which may be as simple as SNPs propagate through their expression in the proteins, their interactions with metabolites or other proteins, until a phenotypic expression. Genotypes resulting in an increased phenotypic fitness under an encompassing environment get selected favorably. ^3,4^ The effect of mutations can however be detrimental as well. Various disorders, such as diabetes or cancers, ^5,6^ and public health concerns such as antibiotic resistance can be traced back to genetic mutations. ^7–9^ Understanding how the effects of distal mutations propagate within the same enzyme is a challenge on its own. Predicting the change in organismal fitness upon amino acid point mutations in proteins is further complicated, since fitness is a downstream effect and an immediate correlation with changes in structural stability and dynamics of the protein may not be easy. However, such an understanding will have an enormous impact, whether it is for identifying disease causing mutations in human genome or for designing drugs against unicellular bacteria.

Mutational landscape analyses had been resource demanding, involving mutations of the genes encoding the protein, their expression, purification and characterization of the effects *in vitro* or *in cellulo*. Despite this difficulty, hundreds of single point mutations, site directed or random mutations, or even systematic alanine scan mutagenesis ^10,11^ experiments were performed on many interesting proteins. ^12^ The effects of mutations on protein folding, stability, enzymatic activity, etc, have been classified as beneficial, neutral, and detrimental to the function of the proteins. Valuable sequence-function relationships can be constructed for the enzyme, with mutational robustness as well as functionally important hotspot residues. However, development of high-throughput technologies, and an interest in protein engineering have driven newer developments in the exploration of mutational landscapes. ^13–15^ Notable among them is the Deep Mutational Scanning methodology, which can perform *>* 10^5^ mutations in a series of experiments.

Deep mutational scanning experiments explore the functional phenotypic effects of thousands of mutants by way of massive sequencing. ^16^ Coupled with a high throughput screening, this approach allows the quantification of the phenotypic changes resulting from the mutations. In principle, the methodology is about an extensive and exhaustive single point mutational scan, where every single amino acid in the protein is replaced with all 19 alternative possibilities. Such detailed studies which identify the enrichment or decimation of each amino acid mutation by functional role selection provide sufficient rationale for selecting target residues for protein engineering. While it has been clearly demonstrated that such deep scan can in principle be performed, the number of such studies is way too less compared to the number of identified protein sequences. The success of the method, its potential for protein engineering, necessitates that alternative ways of predicting the fitness changes to all possible amino acid mutations be developed. Here comes the importance of computational methods. The computational methods using structure and sequence have been used for understanding the function of proteins, in rational design of drugs. *In silico* methods have been applied successfully in several areas of computational biology, from molecular mechanisms of protein function using atomistic simulations to quantitative structure activity relationships using physico-chemical properties of drugs to predict the downstream effects of the drug such as the survival rate of bacteria.

Several computational approaches which use information of the protein sequence, its homologs and/or structure have been developed to score the relative importance of the mutations. SIFT ^17^ uses the amino acid frequencies at each position in a mulitple sequence alignment of homologous proteins to predict whether an amino acid substitution at a particular position in a protein will be neutral or detrimental to its phenotypes. SNAP^18^ is also a sequence information based tool and uses machine learning approach for predicting the effect of mutations. Other models using three-dimensional structural information of the proteins, such as SNPs3d ^19^ are trained on disease and non-disease related data using Support Vector Machine methods to recognize the structural patterns for mutational intolerance. Hybrid models, such as PolyPhen, ^20^ combine available sequence and structural data information into rule based systems to improve the classification of the mutations to potentially non-neutral and neutral. Very recently an unsupervised method based on evolutionary statistical energies computed from sequence covariation ^21^(EVmutation) for the prediction of effects of point mutations was proposed.

All these models except EVmutation act as classifiers, and none have the flexibility to adapt when the phenotypic selection criterion is changed. Predicting the downstream effects of any mutation under any external selection pressure is not easy. While it may be too soon for computational methods to completely replace wet-lab experiments, at this stage they may be helpful in reducing the number of experiments required, by prioritizing them rationally. One can combine a partial deep mutation scan under the desired phenotypic screen with the computational models to achieve the desired predictions. The next step in the evolution of the models is thus a combination of the partial data from deep mutational scan, with sequence, co-evolution and structural information. This is the direction we explore in Deep2Full, which aims at developing a new paradigm of quantitatively predicting the functional outcomes of a full mutational scan by using minimal information from experiments.

## 2 Results and Discussion

### 2.1 Choice of parameters

To computationally simulate the effect of deep mutational scan we combined information from structure, sequence and co-evolution. Seventeen different variables from structural and sequence information of the protein were used. The *structural factors* for each amino acid that could be calculated from a reference protein structure were included in the model - (1) Solvent accessible surface area (SASA) (2) Secondary structural order, with a binary value 1 if the residue is part of a α-helix or β-sheet, and 0 otherwise (3) Number of structural contacts an amino acid has with a 4Å cutoff (4) Average commute time, ^22^ which reflects the average connectivity of a given amino acid with the rest of the protein. The second group of independent parameters were based on the *sequence information* - (5) BLOSUM substitution matrix, which is the probability of substitution of the residue in the wild type by other residues is inferred from evolutionary information ^23^ (6) Hydrophobicity on the Kyte-Dolittle scale ^24^ of the amino acid after mutation (7) Hydrophobicity of the amino acid in the wild type (8) Position specific substitution matrix (PSSM) score for the amino acid after mutation, obtained from PSI-BLAST and (9) PSSM score for the wild-type amino acid (10) Conservation of the amino acid. The third group of independent parameters were based on the properties of *co-evolutionary networks* that were constructed using the multiple sequence alignment (MSA) of hundreds of homologous proteins (**Methods** section). This group which is supposed to reflect the importance of an amino acid in an undirected co-evolutionary network - (11) Average co-evolution correlation of each amino acid (12) Degree centrality, the number of nodes to which a node is connected (13) Betweenness centrality, quantifying the importance of a node in connecting other pairs of residues (14) Closeness centrality, the inverse of the sum of distances to all other nodes (15) Eigenvector centrality, which considers not just the number of connections a node has, but also the connectivity of the immediately connected nodes. Further directed network information (**Methods** section) are included into the model - (16) Impact factor, the number of compensatory mutations required for mutation at residue is calculated based on conditional probabilities (17) Dependency factor, which is the counterpart of impact factor is the number of residues which are likely to influence a mutation in a given protein.

### 2.2 Neural network model

Although the scope of the model is general, in this work, the model is illustrated by complementing the fitness changes in *E. coli* with single point mutations in TEM-1 β-lactamase when exposed to ampicillin. ^25^ A part of the quantitative cellular fitness data obtained from the deep mutational scan of TEM-1 β-lactamase was used for training and validation, and the rest is used for testing the predictions. β-lactamase provides resistance to penicillin-class beta-lactum antibiotics by hydrolyzing the antibiotics. So for quantifying the effect of mutation on function, the selection criterion used was survival under ampicillin presence. The experimental data consists of relative fitness of the mutants with respect to the wild type under different ampicillin concentrations. Neural network method is similar in philosophy to the goal of predicting the downstream effects of the mutations, as the method uses an *unspecified* model which allows inputs and predicts the outputs. The clarity of what happens at the intermediate stages, also known as layers, is compromised in favor of the end results it generates. While it lacks the simplicity of a linear regression model, it can in principle embody all the complex nonlinear interactions that occur at the different stages of the effect propagation, starting from the mutation and ending with the change in fitness. A feedforward neural network with Levenberg-Marquardt back-propagation was used with the NNM module of Matlab. A single hidden layer is used for prediction. The number of hidden neurons is varied and the optimum is decided based on the mean squared error (MSE) calculated over all data points. For a fixed number of hidden neurons, multiple models were created with random initializations for the weights and biases. With every chosen data set, neural network model was built by dividing the data into training, validation and prediction sets. It may seem restrictive that the training and test sets are based on the data from β-lactamase. However, the goal of Deep2Full is to use the context dependent experimental data generated using a phenotypic selection criterion, to rationally reduce the number of such experiments. The neural network model is used for exploring quantitative predictions of fitness when several fractions of the full mutational data is available by way of deep mutational scan.

### 2.3 Full mutational scan

Initially, a neural network model was built using 85% of the complete experimental data obtained at 2500 µ*g*/ml drug concentration, 70% was used for training, 15% for validation and the rest for predictions. Although the available data was obtained from a systematic substitution of every amino acid with all possible choices, the training set we used was based on a random selection of the 70%. The loss or gain in fitness calculated for all possible mutations, including training and predictions are shown in Figure 1. Specifically, a comparison between the experimental observations and the predicted values are shown in Figure 2. With the 17 variables used 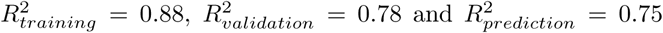 were obtained. Unlike models like SNAP and EVMutation, Deep2Full was trained on the partial data from the experiments to be modeled. While this may seem restrictive, this is a way of factoring in the cellular response under the diverse stressor conditions, concentrations or analogs of antibiotics which may challenge the same enzyme. Deep2Full is thus customized by training on a partial experimental data set of phenotypic selection criteria.

**Figure 1:**
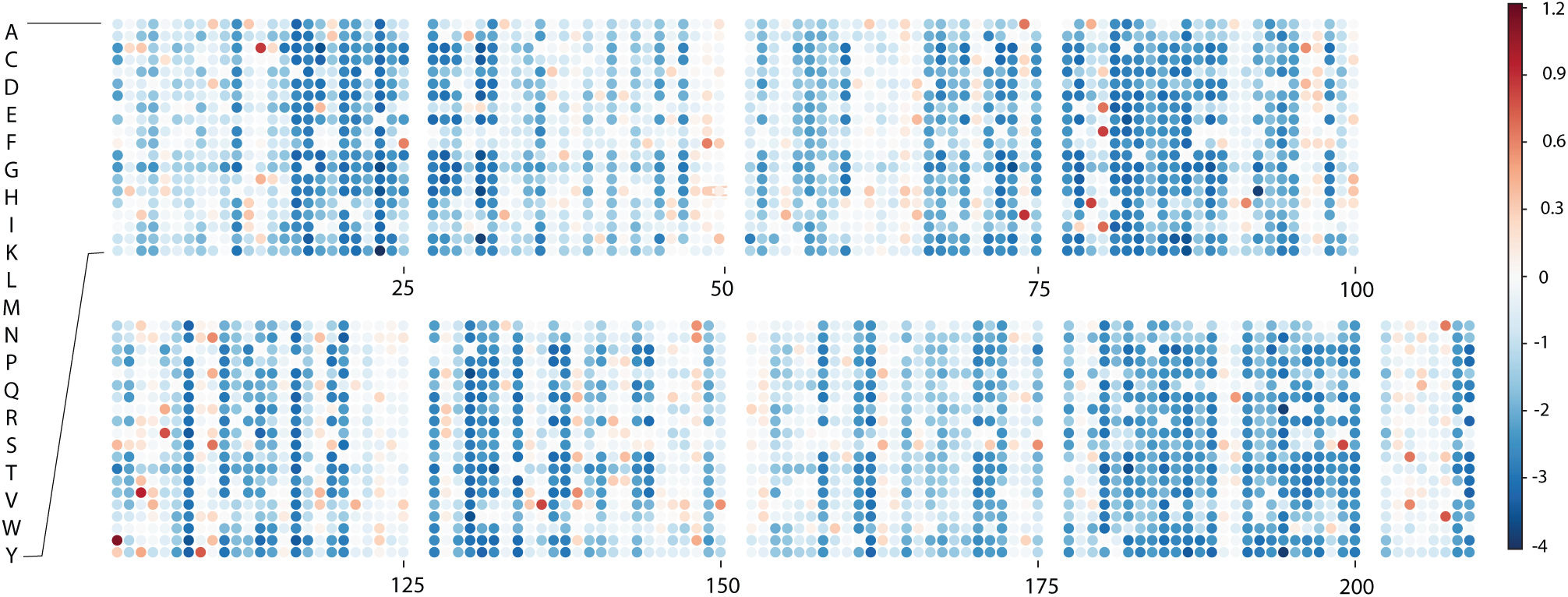
Computational predictions of the quantitative gain or loss in fitness resulting from amino acid changes in β-lactamase. The results, where the change in fitness when every amino acid is changed to all possible alternatives, are an attempt to quantitative deep scan analysis using neural network based computational methods. The complete result set includes the training of the model with experimental data ^25^ when *E. coli* is challenged with 2500 µg/ml ampicilin, validation and test sets. The amino acid numbering according to the reference pdb (1M40) is given in Supplementary Table 1.

**Figure 2:**
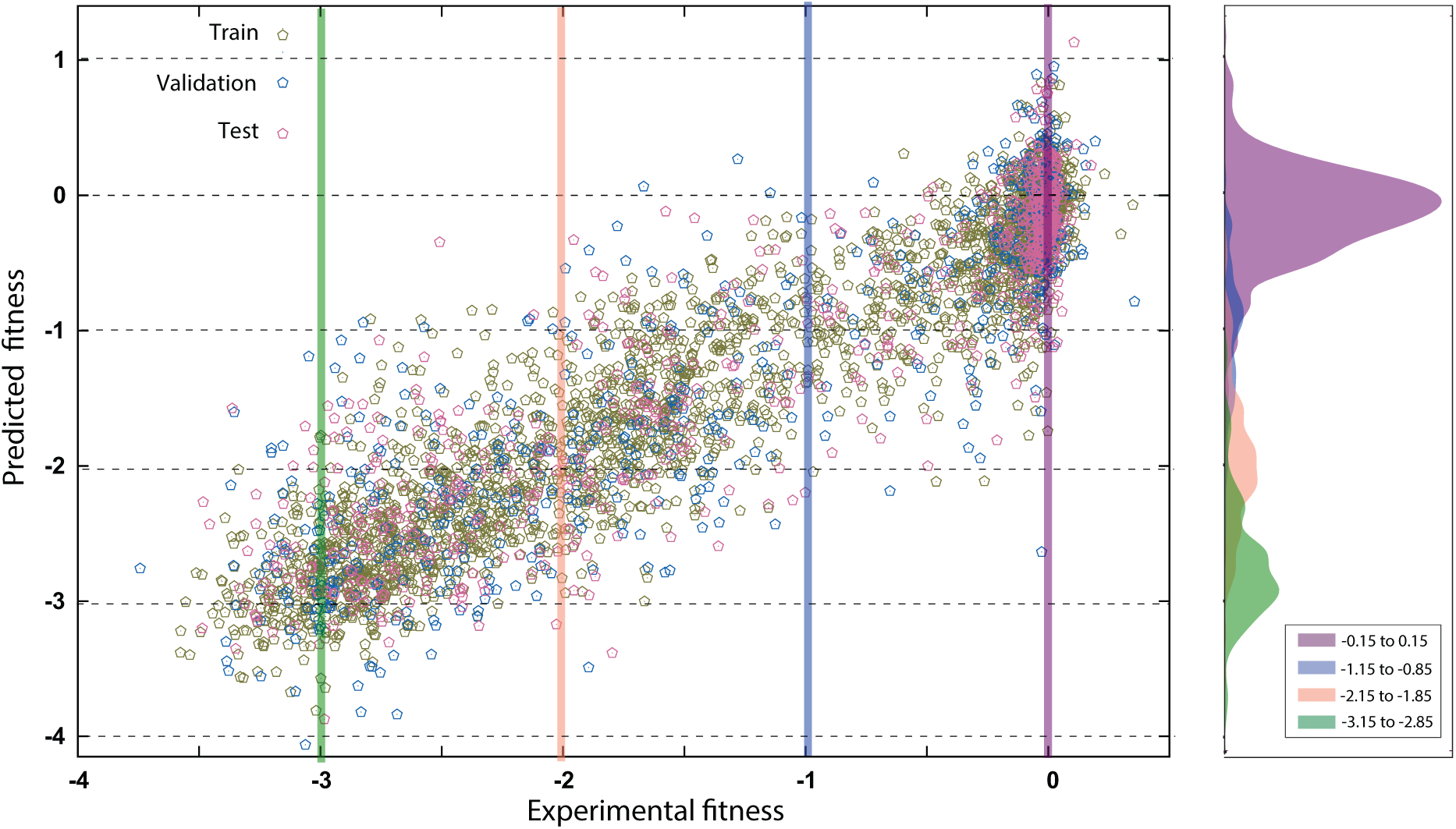
Comparison of the computational results from the neural network model and the experimentally observed fitness. The histogram on the right-hand side shows the distribution of the predicted outcomes, when the experimentally expected fitness values are around −3, −2, −1 and 0.

### 2.4 Effect of individual variable

For investigating the influence of individual input variable on prediction, the input variable is kept fixed at its mean value for all the samples and the network is retrained. This is equivalent to removing the contribution of that particular variable in prediction. Hence the change in mean squared error(MSE) on removal of a variable quantifies the importance of that input parameter. Figure 3 shows the difference in MSE when each variable is replaced by its mean value. BLOSUM which represents the substitution effects based on evolutionary data has the highest contribution to the predictions. Hydrophobicity index of the amino acid to which the mutation is made and the average commute time are the other variables with two significantly higher contributions. In addition to the 17 variables, we also added the statistical coupling energy ^21^ as an additional variable to see if it improved the correlation between the predictions and the observations. No improvement was noticed, possibly because the co-evolutionary effects represented by variables 11 to 17 already implicitly accounted for this factor. So statistical coupling energy was not used in any other analysis in this work. We also analyzed the contributions at a coarse level, creating neural network model for alanine scan mutations using only (1) sequence based variables and (2) structure based variables. The sequence based model is slightly better than the structure based one, 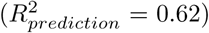 values being 0.55 and 0.44 respectively for the sequence and structure based models.

**Figure 3:**
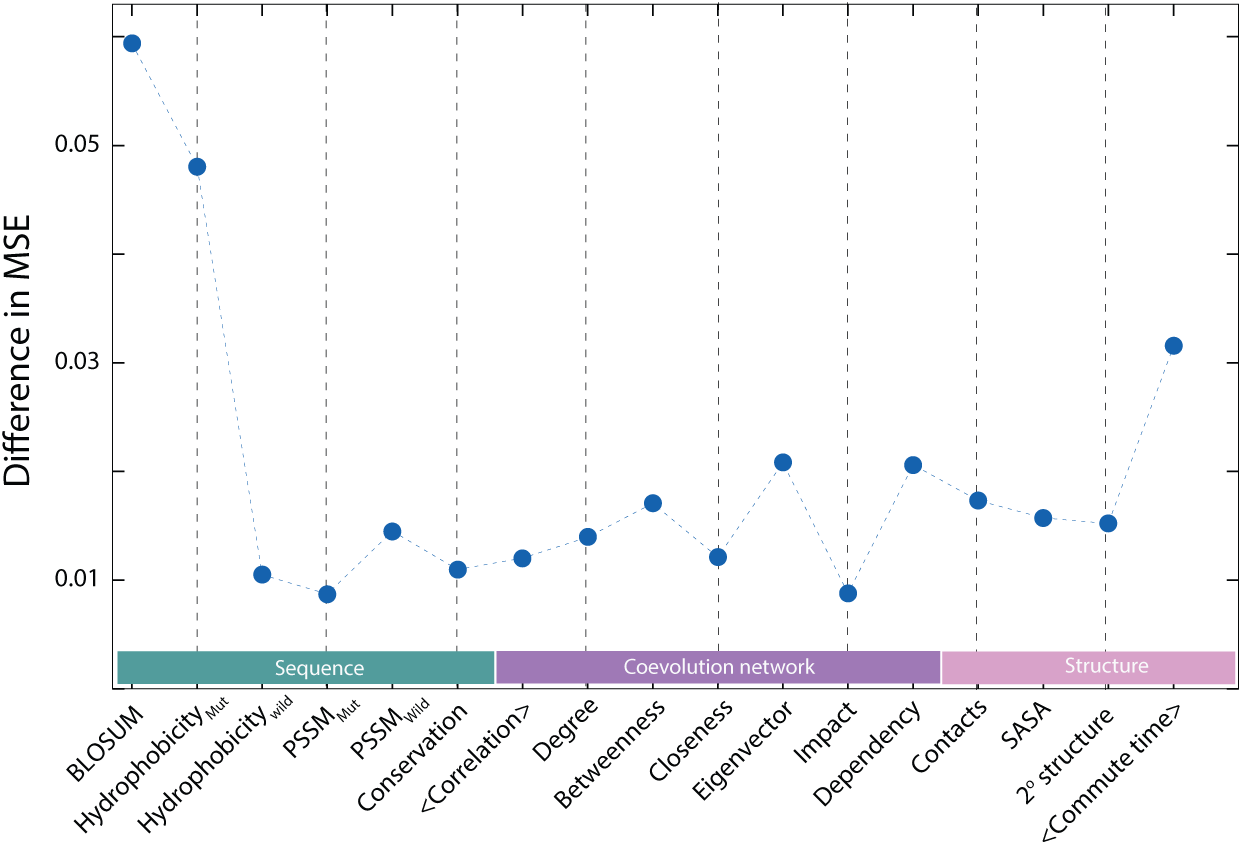
Variable impact analysis for the seventeen variables used in the model is shown. The variable impact is calculated by sequentially replacing each variable by its average value and checking for the change in the mean squared error between the expected and predicted values.

### 2.5 Complementary ANH scan

Alanine scan mutagenesis is a popular biochemical technique which aims to replace all potentially interesting amino acids with alanine, and study the effects on the stability and function of the protein. The choice of alanine is motivated by the goal for standardization and simplicity of the substituted amino acid. Using the experimental data of the fitness response to alanine scans, the outcomes for all other 19 mutational scans were predicted using Deep2Full. However, in our search for the minimal data set which can be useful for understanding the effect of all mutations, alanine proved insufficient as the fitness changes with mutations to all other 19 amino acids could not be predicted satisfactorily (Supplementary Figure 1). It was recently discovered by curating the data from all the deep mutational scan data that the potential impact of an amino acid is understood by scanning with mutations to two amino acids - asparagine (N) and histidine (H). ^26^ Taking cue from this observation, we combined the commonly used alanine (A) scan, with asparagine (N) and histidine (H) scans, thus choosing one from each charge type - hydrophobic (A), polar (N) and charged (H). We then used ANH-scan as a strategy for training the neural network model. The fitness outcomes from ANH scan, which contributed to 3/20 or 15% of the full mutational scan, were used to predict all the remaining 17 amino acid scan results. The minimal ANH scan data was further divided as 85% for training and 15% for validation of the model (Figure 4A). This model was then tested on the predictions for the scan with all 17 other amino acid scans. While the number of data points used is the proportion 3 out of 20 (15%), the results for fitness predictions were in fairly good agreement with the experimental data 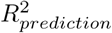 = 0.49, as shown in Figure 4B.

**Figure 4:**
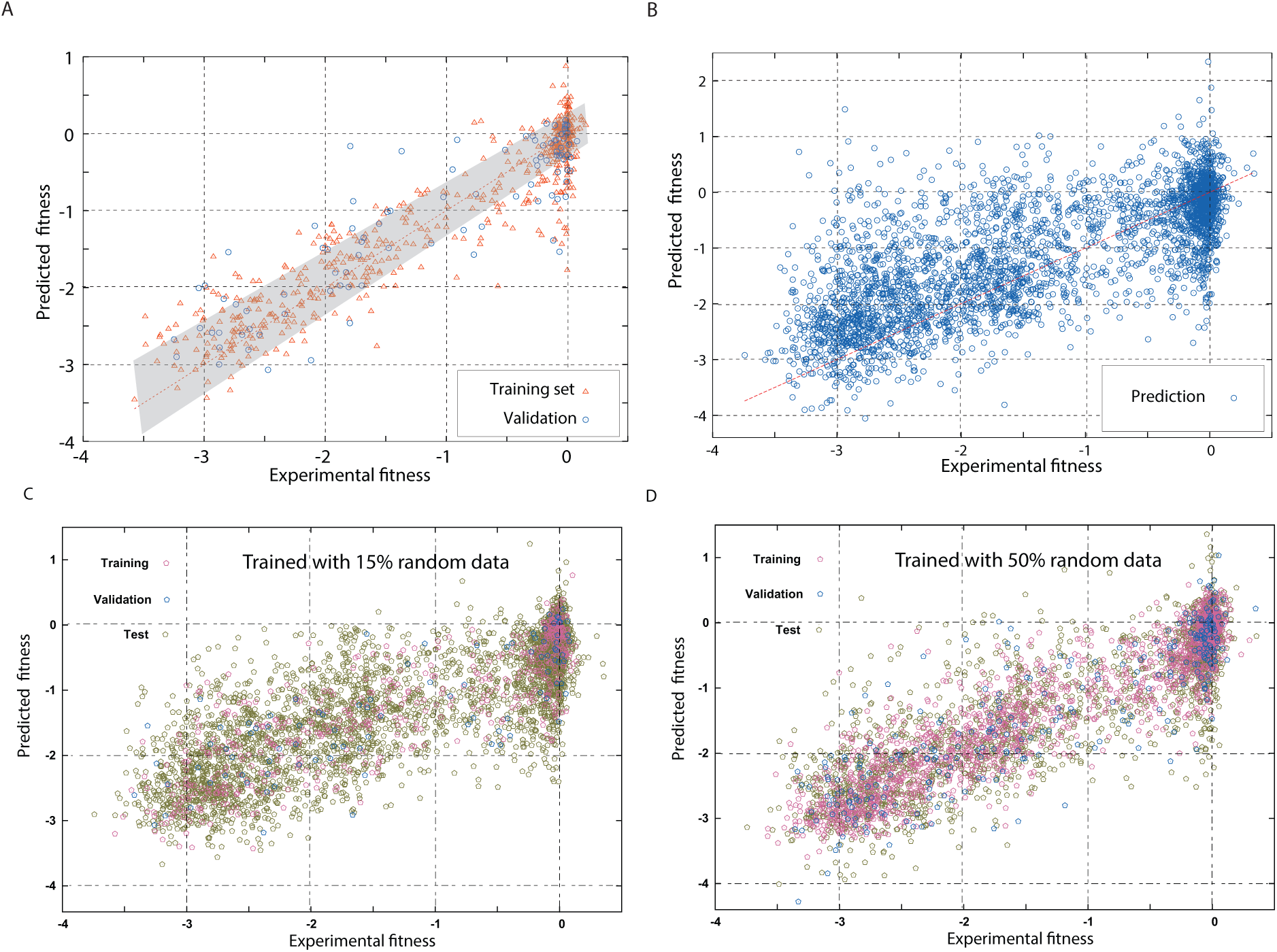
In an extension of the concept of alanine-scan, the fitness outcomes from alanine(A), asparagine(N) and histidine(H)-scans at each amino acid position were used as the training set, and the results for all other 17 mutations at every site were predicted. The graph shows the results from these (A) training and validation (B) prediction. The grey bar in (A) indicates the variation in fitness between different trials of the experiment. The results are compared with using (C) 15% and (D) 50% randomly picked mutations for training, which show that at the same level of training (15%), choosing a random set of mutations is better than a systematic ANH scan.

### 2.6 Random vs. ANH scan

When reasonable fitness outcomes could be predicted with as few as 15% of the full mutational scan data, a question arises whether is the better way to perform the scan. We make a comparison with the outputs when the same few data points are chosen randomly instead of from ANH scan. The motivation for the training the model with the random mutations is that if it indeed is at least as effective as ANH scan, the experimental constraints on a very specific scan will be lesser. Interestingly, we see that when 15% of the full mutational scan data is chosen for a combination of training and validation, the predictions are slightly better than from the predictions of the complementary ANH scan as seen in Figure 4C 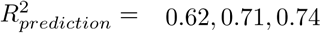 Also in general the distribution of *R*^2^ for each of the amino acids before or after mutation is better with a random choice (Supplementary Figure 4). Of course, increasing the training and validation set further, to 50% makes the predictions better (Figure 4D) and almost comparable to the results from using 85% data in Figure 2.

### 2.7 Comparison with scoring models

Scoring models such as SNAP have been used for classifying the mutations as fitness-neutral or non-neutral.^18^ Other recent models have shown a good correlation between the evolutionary statistical energy ^21^, the fitness score obtained from co-evolutionary model and the fitness variations observed in deep mutational scan. Unlike Deep2Full, these models are unsupervised and do not require a partial set of experiments to generate the scores. However, if the goal is to build a model which quantitatively predicts the fitness variations of mutations with minimal wet-lab experiments, the priorities differ. For example, the magnitude of effect of the same β- lactamase mutation in *E. coli* is different at different ampicillin concentrations. Deep2Full focuses on using partial data from mutational scan experiments to customize the model to simultaneous variations in mutational space as well as to the changes in concentrations of stressors. A similar approach using SNAP or EVM scores would also require training the unsupervised computational scores against the experimentally observed fitness variations to develop a model, possibly a linear regression model relating the score with the fitness. As shown in Supplementary Figure 5, for the specific example considered, the relations between the scores and cellular fitness using 15% and 50% of randomly selected mutations are at least as noisy or worse than the models we used.

**Figure 5:**
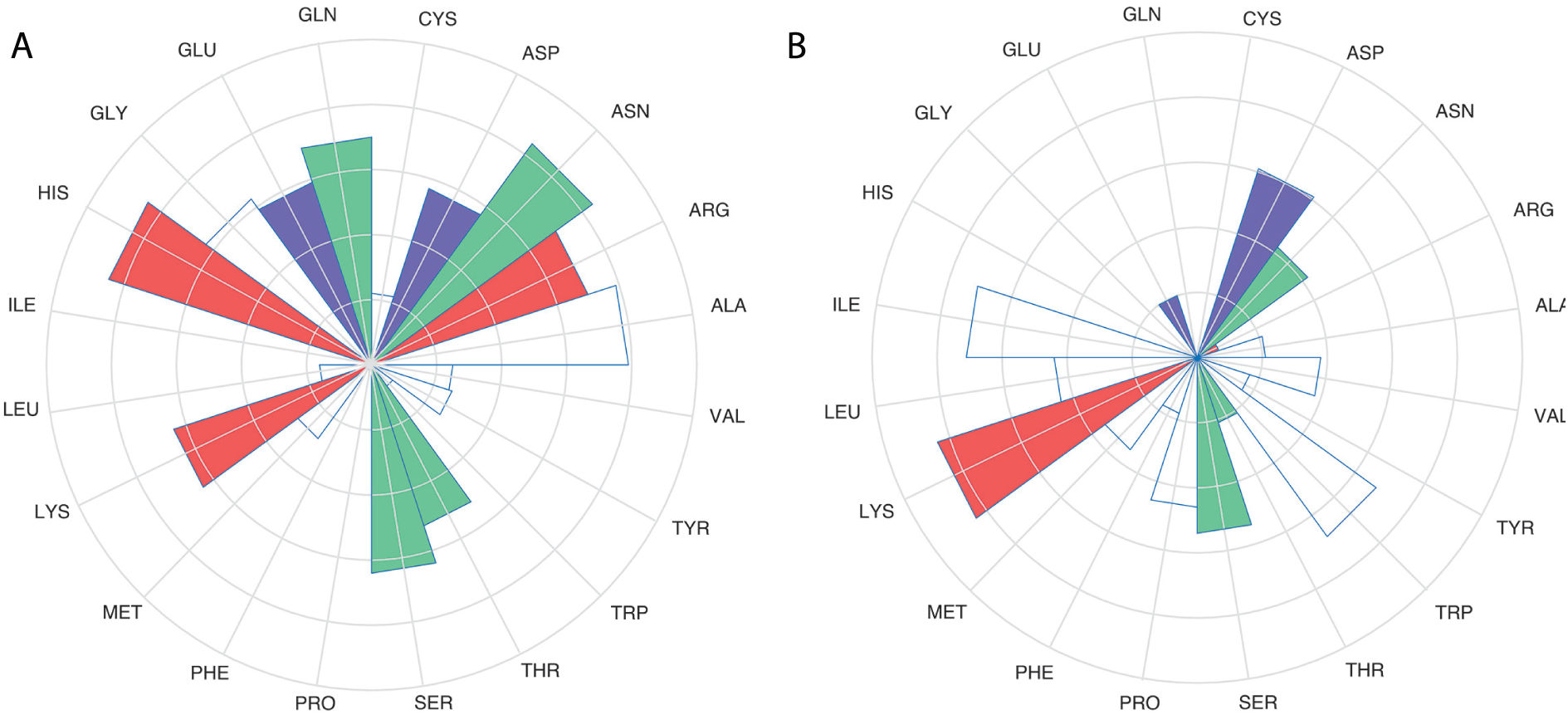
Computational deep mutational scan predictions using ANH scan data for training were sorted using (A) the amino acid after mutation (B) the amino acid in the wild type. Amino acids are colored according to their type: red (positive), blue (negative), green (polar), white (hydrophobic)

### 2.8 Dose-response

Deep mutational scan experiments are in general repeated for several assay conditions, specifically the fitness changes at different concentrations of the drug were measured for *E. coli*. ^25^ The question we explore here is whether the predictive computational models can be useful for reducing the number of experimental trials. The fitness-data corresponding to all 4997 mutations of β-lactamase at six different concentrations was clustered to see the different groups of amino acid mutations which show similar early or late response patterns (Figure 6A) according to the drug concentration. The patterns among the bundles of amino acid mutations which behave similarly or cluster together are then shown as a function of concentration (Figure 6B). It can be seen that these mutations do follow a regular dose-response pattern, a sigmoidal response curve, with log [ampicillin]. Thus the same paradigm can be extended: that by knowing partial data at different concentrations, which will be helpful in developing the dose-response curves, the fitness variations arising from every single amino acid mutation can be predicted. However, considering that the differences that were seen between the two different trials of the same experiment, we do not attempt the quantitative concentration dependent full mutational scan at this stage.

**Figure 6:**
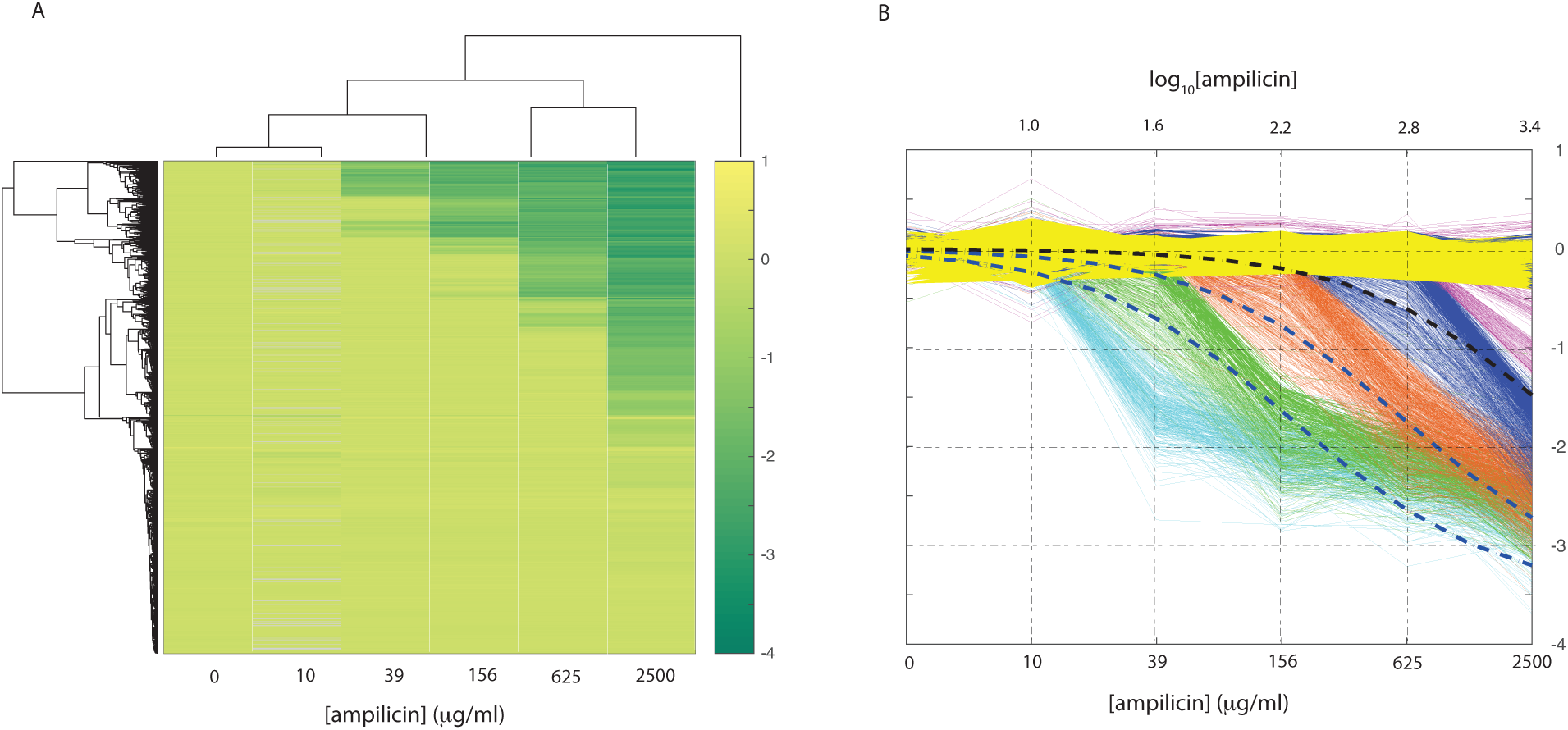
The experimental data of about 4997 mutations of β-lactamase was clustered to see the different dose-response groups. (A) Hierarchically clustered data of the fitness response (B) Bundles of data representing the different clusters which have comparable dose response curves. The dashed lines are fits to sigmoid curves of fitness clusters with log([ampicillin]).

### 2.9 Prediction Quality Analysis

The quality of predictions were judged from three different measures - (1) the overall 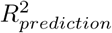, which was good even when 15% of the data is used. The quality improves as more data is used for training, 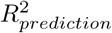 = 0.62,0.71,0.74 corresponding to 15%, 50% and 85% combined training and validation data. (2) the prediction data was segregated either based on the amino acid before or after mutation. The outcomes for some of the amino acids are relatively poor as seen from the individual regression plots (Supplementary Figures 2 and 3), which are summarized using their 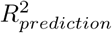 values in Figure 5. It can be noticed that the effects of some amino acid mutations do not span the entire range, hence predictions could not be improved. (3) For an expected experimental fitness, the variation in the predicted values. This is summarized in Figure 2 with histograms of predicted fitness generated from the expected fitness variation around −3, −2, −1 and 0. While these histograms show a variation relative to the expected fitness, it must be noted that even in different trials of the experiment, there is a significant variation. This variation in the predictions is comparable to the variation in the observed fitness in different experimental trials that is marked in Figure 4A.

## 3 Conclusions

Deep2Full is developed as a new paradigm for developing computational models customized with deep mutational scan data from some concentrations to quantitatively predict the fitness outcomes of a full-set of mutations and at multiple concentrations of the stressors. Neural network model and seventeen structural, sequence and co-evolutionary variables were used to make these predictions. By combining this phenotypic deep scan data with structure, sequence and co-evolutionary information, reliable predictions of the full set of deep mutational scan was achieved. Model prediction can be improved as the repeatability in the different trials of the experiments improves. Deep2Full is a way of rationally reducing the number of mutagenesis experiments required to make the fitness predictions in a full mutational scan.

## 4 Methods

### Co-evolution network and properties

Multiple Sequence Alignment(MSA) for the β-lactamase family was obtained from Pfam (Pfam ID: PF13354) and sequences with a gap frequency less than 20% were used for the analysis. Consensus sequence was generated using the amino acid that most occurs in a given position. Following the Statistical Coupling Analysis protocol, ^27^ MSA was converted into a boolean sequence, with a 1 if the amino acid is the same as in the consensus sequence and 0 otherwise.

### Undirected network

The co-evolutionary relation between two amino acids *i* and *j* is calculated as proposed by Halabi et. al. ^27^, 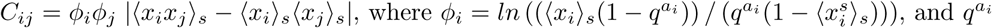 is the probability with which the amino acid *a_i_* at position *i* in the consensus sequence occurs among all proteins. *x*_*i*_ is the *i*^th^ column in the boolean sequence and <>_s_ denotes the average over sequences. The co-evolutionary matrix is converted into a network representation using a cutoff *c*. If *C*_*ij*_ > *c*, we consider an undirected co-evolutionary network *i* — *j* to be present. In the present analysis weighted co-evolutionary matrix was used and the cut-off chosen was 1.5. Using the network of amino acids is built using this criterion, different centrality measures were calculated -eigen vector centrality, degree, etc.

*Directed network*: Using the boolean sequence that was created using multiple sequence alignment, we created a directed influence network, in a co-evolutionary sense, with the following conditional probabilities:

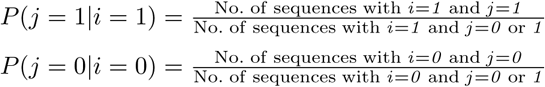

We used a cut-off *P* = 0.87. If both *P* (*j* = 1|*i* = 1) and *P* (*j* = 0|*i* = 0) are simultaneously greater than a value *P*. In this directed network, the number of outgoing links is considered the impact of an amino acid, and the number of incoming links is considered its dependency. The impact and dependence are supposed to summarize how many simultaneous mutations are forced or forced-upon by a mutation.

### Average commute time

The hypothesis that the structural and dynamical connectivity of an amino acid to other amino acids determines the importance of an amino acid has been put forward. ^22^ The average commute time has been used for identifying hotspot amino acids. The resistance matrix is constructed using the number of atom-atom contacts between amino acids *i* and *j*, which are within 4Å. Structure obtained from PDB (1M40) was used for this definition. The resistance matrix is used for average commute time calculations as per the algorithm suggested in Ref. ^22^

## Acknowledgement

We thank Prof. Hemalatha Balaram for the helpful discussions.

## Supplementary information

**Figure 1:**
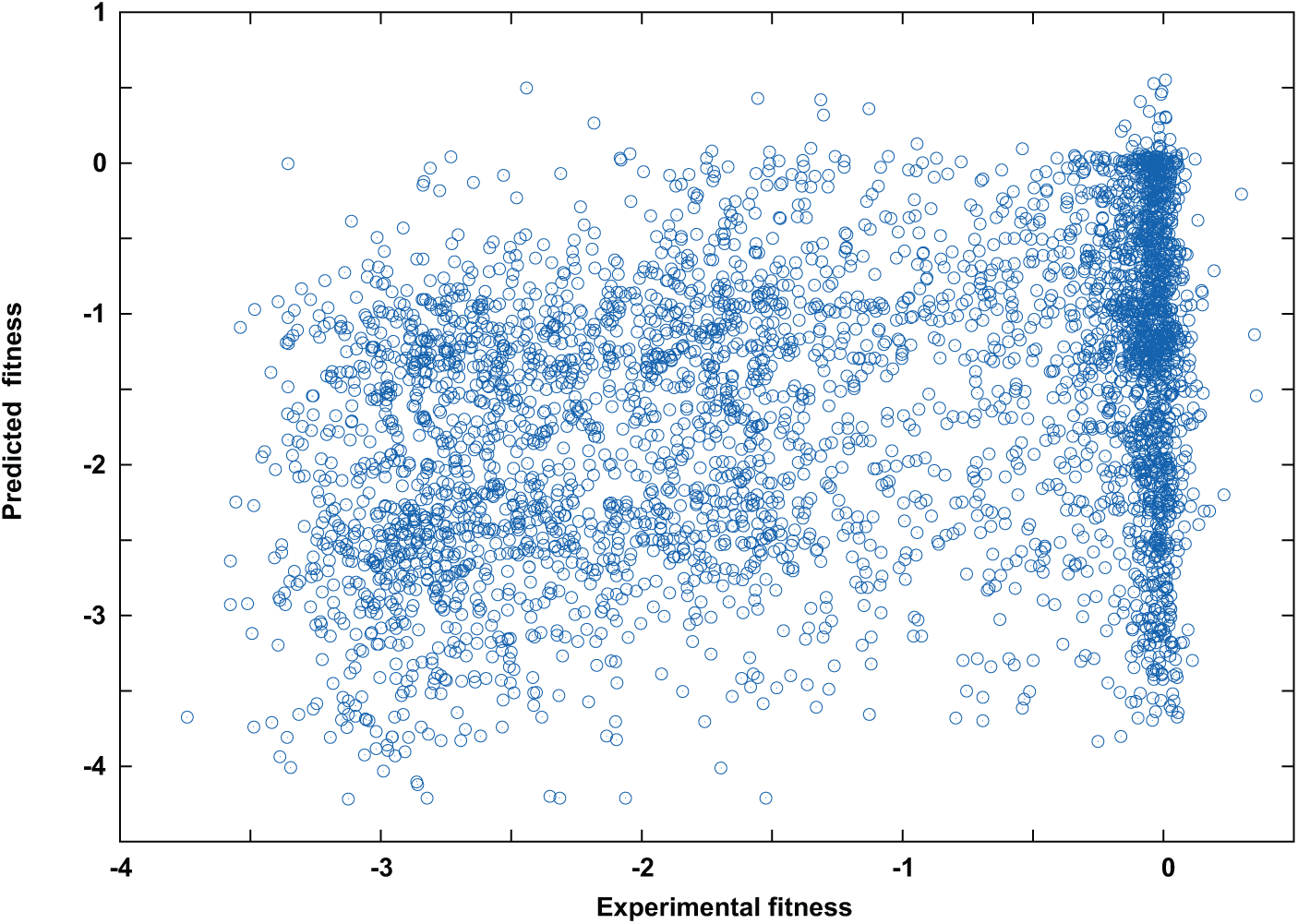
Alanine scan data of the β-lactamase was used to predict the phenotypic fitness changes arising from all other 19 mutational scans. The results from using 1/20 mutational scans (5% of the data) for training are seen to be poor.

**Figure 2:**
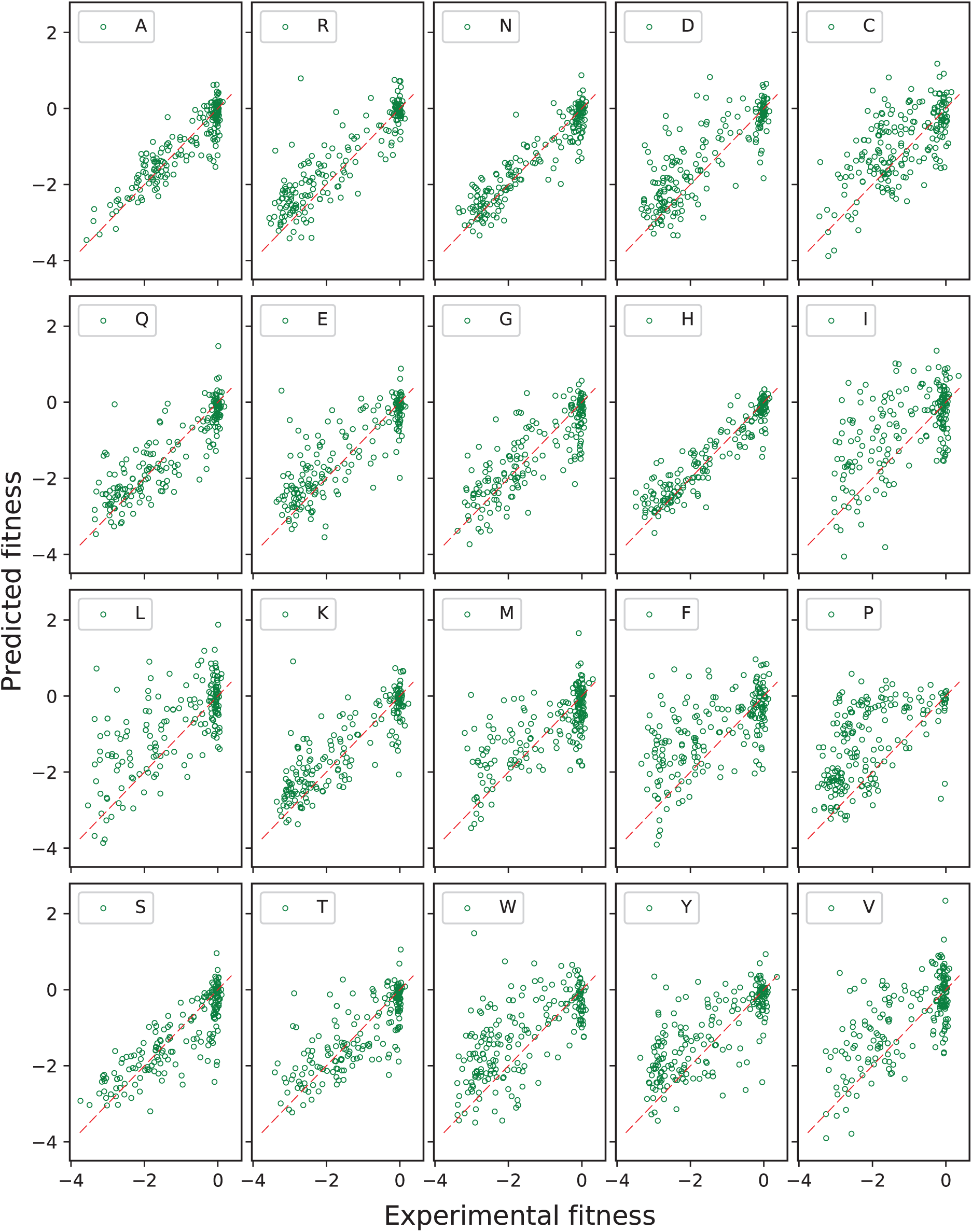
Predictions from ANH were analyzed by classifying them according to the amino acid to which mutation is performed. The analysis shows the predictability for all amino acids is comparable. However, the predictions to alanine (A), asparagine (N), histidine (N) are notably better because of the training set that was used. The dashed red lines are guidelines with slope −1.

**Figure 3:**
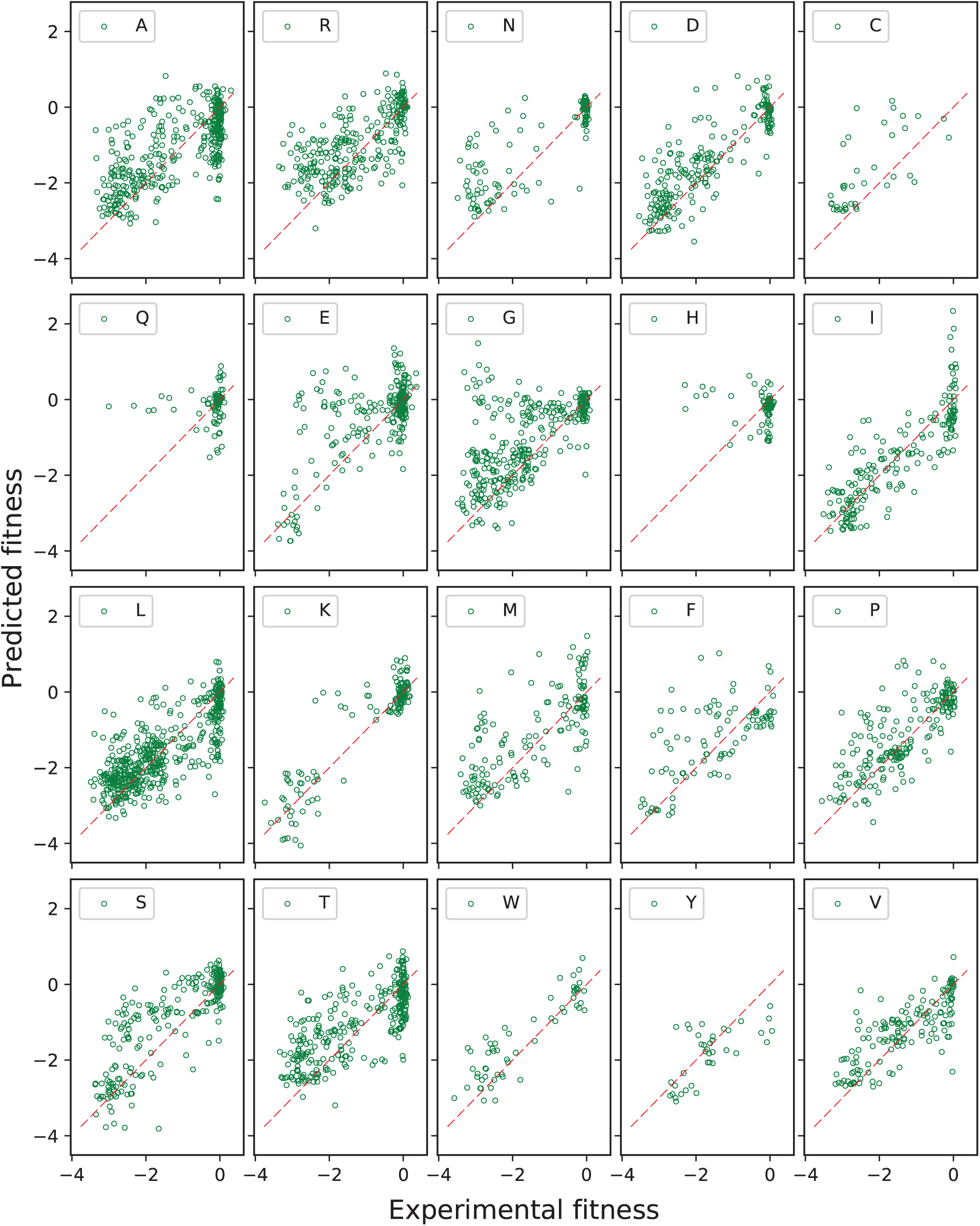
Predictions from ANH were also analyzed by classifying them according to the wild type amino acid from which mutation is performed. The analysis shows the predictability for all amino acids is comparable. The dashed red lines are guidelines with slope −1.

**Figure 4:**
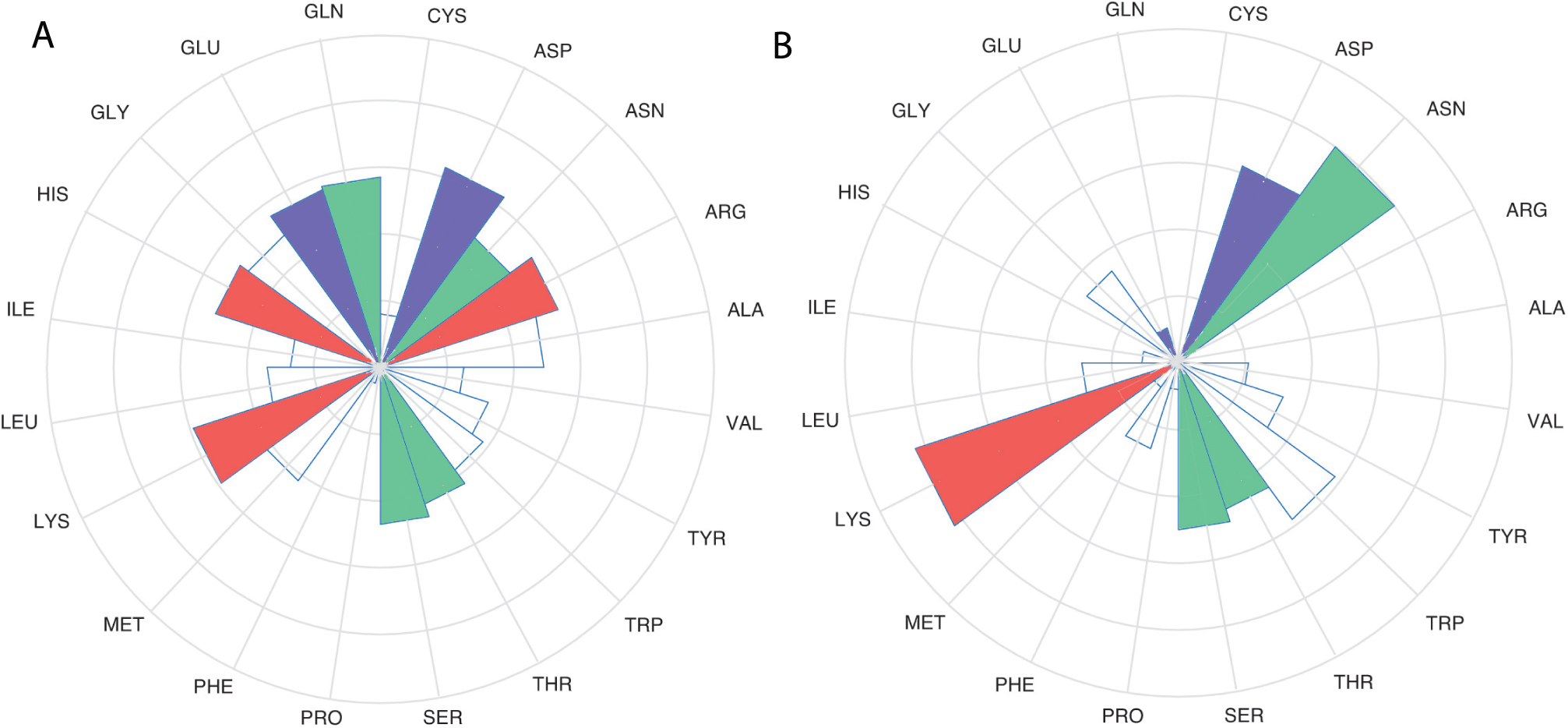
Computational deep mutational scan predictions trained on a random 15% of the experiments. The results were sorted using (A) the amino acid after mutation (B) the starting amino acid. Amino acids are colored according to their type: red (positive), blue (negative), green (polar), white (hydrophobic)

**Figure 5:**
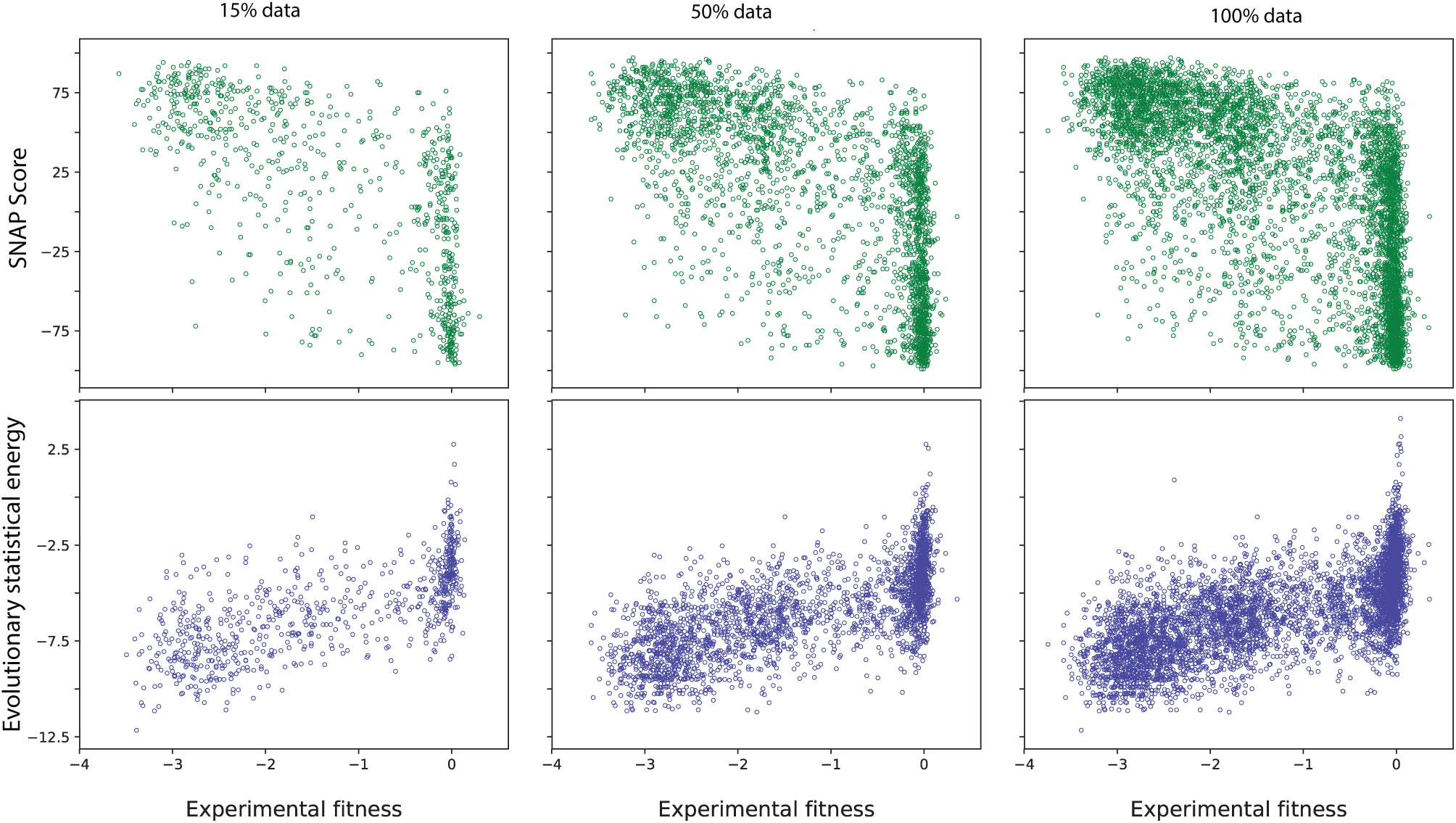
A comparison of how the scores from SNAP and Evolutionary statistical energy relate to the experimentally observed fitness of *E. coli* arising from β-lactmase mutations is shown in this figure. The scores are compared for the complete experimental data as well as by randomly selecting 15%, 50% of the data.

## Supplementary Table 1

**Table 1:** The table indicates the index used in Figure 1 of main article and the amino acid(a. a.) and its numbering according to reference wild type pdb (1M40)

| Sl. No. | a. a. number | a.a | Sl. No. | a. a. number | a.a | Sl. No. | a. a. number | a.a |
| --- | --- | --- | --- | --- | --- | --- | --- | --- |
| 1 | 51 | L | 36 | 86 | A | 71 | 121 | E |
| 2 | 52 | N | 37 | 87 | G | 72 | 122 | L |
| 3 | 53 | S | 38 | 88 | Q | 73 | 123 | C |
| 4 | 54 | G | 39 | 89 | E | 74 | 124 | S |
| 5 | 55 | K | 40 | 90 | Q | 75 | 125 | A |
| 6 | 56 | I | 41 | 91 | L | 76 | 126 | A |
| 7 | 57 | L | 42 | 92 | G | 77 | 127 | I |
| 8 | 58 | E | 43 | 93 | R | 78 | 128 | T |
| 9 | 59 | S | 44 | 94 | R | 79 | 129 | M |
| 10 | 60 | F | 45 | 95 | I | 80 | 130 | S |
| 11 | 61 | R | 46 | 96 | H | 81 | 131 | D |
| 12 | 62 | P | 47 | 97 | Y | 82 | 132 | N |
| 13 | 63 | E | 48 | 98 | S | 83 | 133 | T |
| 14 | 64 | E | 49 | 99 | Q | 84 | 134 | A |
| 15 | 65 | R | 50 | 100 | N | 85 | 135 | A |
| 16 | 66 | F | 51 | 101 | D | 86 | 136 | N |
| 17 | 67 | P | 52 | 102 | L | 87 | 137 | L |
| 18 | 68 | M | 53 | 103 | V | 88 | 138 | L |
| 19 | 69 | M | 54 | 104 | E | 89 | 139 | L |
| 20 | 70 | S | 55 | 105 | Y | 90 | 140 | T |
| 21 | 71 | T | 56 | 106 | S | 91 | 141 | T |
| 22 | 72 | F | 57 | 107 | P | 92 | 142 | I |
| 23 | 73 | K | 58 | 108 | V | 93 | 143 | G |
| 24 | 74 | V | 59 | 109 | T | 94 | 144 | G |
| 25 | 75 | L | 60 | 110 | E | 95 | 145 | P |
| 26 | 76 | L | 61 | 111 | K | 96 | 146 | K |
| 27 | 77 | C | 62 | 112 | H | 97 | 147 | E |
| 28 | 78 | G | 63 | 113 | L | 98 | 148 | L |
| 29 | 79 | A | 64 | 114 | T | 99 | 149 | T |
| 30 | 80 | V | 65 | 115 | D | 100 | 150 | A |
| 31 | 81 | L | 66 | 116 | G | 101 | 151 | F |
| 32 | 82 | S | 67 | 117 | M | 102 | 152 | L |
| 33 | 83 | R | 68 | 118 | T | 103 | 153 | H |
| 34 | 84 | V | 69 | 119 | V | 104 | 154 | N |
| 35 | 85 | D | 70 | 120 | R | 105 | 155 | M |
| 106 | 156 | G | 141 | 191 | R | 176 | 226 | P |
| 107 | 157 | D | 142 | 192 | K | 177 | 227 | A |
| 108 | 158 | H | 143 | 193 | L | 178 | 228 | G |
| 109 | 159 | V | 144 | 194 | L | 179 | 229 | W |
| 110 | 160 | T | 145 | 195 | T | 180 | 230 | F |
| 111 | 161 | R | 146 | 196 | G | 181 | 231 | I |
| 112 | 162 | L | 147 | 197 | E | 182 | 232 | A |
| 113 | 163 | D | 148 | 198 | L | 183 | 233 | D |
| 114 | 164 | R | 149 | 199 | L | 184 | 234 | K |
| 115 | 165 | W | 150 | 200 | T | 185 | 235 | S |
| 116 | 166 | E | 151 | 201 | L | 186 | 236 | G |
| 117 | 167 | P | 152 | 202 | A | 187 | 237 | A |
| 118 | 168 | E | 153 | 203 | S | 188 | 238 | G |
| 119 | 169 | L | 154 | 204 | R | 189 | 240 | E |
| 120 | 170 | N | 155 | 205 | Q | 190 | 241 | R |
| 121 | 171 | E | 156 | 206 | Q | 191 | 242 | G |
| 122 | 172 | A | 157 | 207 | L | 192 | 243 | S |
| 123 | 173 | I | 158 | 208 | I | 193 | 244 | R |
| 124 | 174 | P | 159 | 209 | D | 194 | 245 | G |
| 125 | 175 | N | 160 | 210 | W | 195 | 246 | I |
| 126 | 176 | D | 161 | 211 | M | 196 | 247 | I |
| 127 | 177 | E | 162 | 212 | E | 197 | 248 | A |
| 128 | 178 | R | 163 | 213 | A | 198 | 249 | A |
| 129 | 179 | D | 164 | 214 | D | 199 | 250 | L |
| 130 | 180 | T | 165 | 215 | K | 200 | 251 | G |
| 131 | 181 | T | 166 | 216 | V | 201 | 252 | P |
| 132 | 182 | M | 167 | 217 | A | 202 | 254 | D |
| 133 | 183 | P | 168 | 218 | G | 203 | 255 | G |
| 134 | 184 | A | 169 | 219 | P | 204 | 256 | K |
| 135 | 185 | A | 170 | 220 | L | 205 | 257 | P |
| 136 | 186 | M | 171 | 221 | L | 206 | 258 | S |
| 137 | 187 | A | 172 | 222 | R | 207 | 259 | R |
| 138 | 188 | T | 173 | 223 | S | 208 | 260 | I |
| 139 | 189 | T | 174 | 224 | A |  |  |  |
| 140 | 190 | L | 175 | 225 | L |  |  |  |

## References

[1] M. Nachman, “Single nucleotide polymorphisms and recombination rate in humans,” TRENDS IN GENETICS, vol. 17, pp. 481–485, SEP 2001.

[2] L. B. Barreiro, G. Laval, H. Quach, E. Patin, and L. Quintana-Murci, “Natural selection has driven population differentiation in modern humans,” NATURE GENETICS, vol. 40, pp. 340–345, MAR 2008.

[3] H. Orr, “Fitness and its role in evolutionary genetics,” Nature Reviews Genetics, vol. 10, pp. 531–539, 2009.

[4] H. Ellegren and B. Sheldon, “Genetic basis of fitness differences in natural populations” Nature, vol. 452, pp. 169–175, 2008.

[5] J. Gudmundsson, P. Sulem, V. Steinthorsdottir, J. T. Bergthorsson, G. Thorleifsson, U. Thorsteinsdottir, A. Kong, and K. Stefansson, “Two variants on chromosome 17 confer prostate cancer risk, and the one in TCF2 protects against type 2 diabetes,” Nature Genetics, vol. 39, p. 977–983, 2007.

[6] M. O’Hayre, J. Vazquez-Prado, I. Kufareva, E. W. Stawiski, T. M. Handel, S. Seshagiri, and J. S. Gutkind, “The emerging mutational landscape of G proteins and G-protein-coupled receptors in cancer,” Nature Genetics, vol. 13, pp. 412–424, 2013.

[7] C. Walsh, “Molecular mechanisms that confer antibacterial drug resistance,” Nature, vol. 406, pp. 775–781, 2000.

[8] E. D. Brown and G. D. Wright, “Antibacterial drug discovery in the resistance era,” Nature, vol. 529, p. 336–343, 2017.

[9] M. O. A. Sommer, C. Munck, R. V. Toft-Kehler, and D. I. Andersson, “Molecular mechanisms that confer antibacterialdrugresistance,” Nature, vol. 406, pp. 775–781, 2000.

[10] B. Cunningham and J. Wells, “High-resolution epitope mapping of high-receptor interactions by alanine-scanningmutagenesis,” Science, vol. 244, pp. 1081–1085, JUN 2 1989.

[11] T. Kortemme, D. E. Kim, and D. Baker, “Computational alanine scanning of protein-protein interfaces,” Science STKE, vol., p. pl2, 2004.

[12] P. A. Romero and F. H. Arnold, “Exploring protein fitness landscapes by directed evolution,” Nature Reviews Molecular Cell Biology, vol. 10, pp. 866–876, 2009.

[13] D. M. Fowler, C. L. Araya, S. J. Fleishman, E. H. Kellogg, J. J. Stephany, D. Baker, and S. Fields, “High-resolution mapping of protein sequence-function relationships,” NATURE METHODS, vol. 7, pp. 741–U108, SEP 2010.

[14] R. T. Hietpas, J. D. Jensen, and D. N. A. Bolon, “Experimental illumination of a fitness landscape,” Proceedings of the National Academy of Sciences of the United States of America, vol. 108, pp. 7896–7901, MAY 10 2011.

[15] L. Zheng, U. Baumann, and J.-L. Reymond, “An efficient one-step site-directed and site-saturation mutagenesis protocol,” Nucleic Acids Research, vol. 32, p. e115, 2004.

[16] C. L. Araya and D. M. Fowler, “Deep mutational scanning: assessing protein function on a massive scale,” TRENDS IN BIOTECHNOLOGY, vol. 29, pp. 435–442, SEP 2011.

[17] N.-L. Sim, P. Kumar, J. Hu, S. Henikoff, G. Schneider, and P. C. Ng, “SIFT web server: predicting effects of amino acid substitutions on proteins,” NUCLEIC ACIDS RESEARCH, vol. 40, pp. W452–W457, JUL 2012.

[18] Y. Bromberg, G. Yachdav, and B. Rost, “SNAP predicts effect of mutations on protein function,” BIOINFORMATICS, vol. 24, pp. 2397–2398, OCT 15 2008.

[19] P. Yue, Z. Li, and J. Moult, “Loss of protein structure stability as a major causative factor in monogenic disease,” JOURNAL OF MOLECULAR BIOLOGY, vol. 353, pp. 459–473, OCT 21 2005.

[20] I. A. Adzhubei, S. Schmidt, L. Peshkin, V. E. Ramensky, A. Gerasimova, P. Bork, A. S. Kondrashov, and S. R. Sunyaev, “Amethod and server for predicting damaging missense mutations,” NATURE METHODS, vol. 7, pp. 248–249, APR 2010.

[21] T. A. Hopf, J. B. Ingraham, F. J. Poelwijk, C. P. I. Scharfe, M. Springer, C. Sander, and D. S. Marks, “Mutatione effects predicted from sequence co-variation,” NATURE BIOTECHNOLOGY, vol. 35, pp. 128–135, FEB 2017.

[22] C. Chennubhotla and I. Bahar, “Signal propagation in proteins and relation to equilibrium fluctuations,” PLOS COMPUTATIONAL BIOLOGY, vol. 3, pp. 1716–1726, SEP 2007.

[23] S. Henikoff and J. Henikoff, “AMINO-ACID SUBSTITUTION MATRICES FROM PROTEIN BLOCKS,” PROCEEDINGS OF THE NATIONAL ACADEMY OF SCIENCES OF THE UNITED STATES OF AMERICA, vol. 89, pp. 10915–10919, NOV 15 1992.

[24] J. Kyte and R. Doolittle, “Amino acid scale: Hydropathicityl,” Journal of Molecular Biology, vol. 157, pp. 105–132, 1982.

[25] M. A. Stiffler, D. R. Hekstra, and R. Ranganathan, “Evolvability as a Function of Purifying Selection in TEM-1 beta-Lactamase,” CELL, vol. 160, pp. 882–892, FEB 26 2015.

[26] V. E. Gray, R. J. Hause, and D. M. Fowler, “Analysis of Large-Scale Mutagenesis Data To Assess the Impact of Single Amino Acid Substitutions,” GENETICS, vol. 207, pp. 53–61, SEP 2017.

[27] N. Halabi, O. Rivoire, S. Leibler, and R. Ranganathan, “Protein sectors: evolutionary units of three-dimensional structure,” Cell, vol. 138, no. 4, pp. 774–786, 2009.

